# Chlorotoxin Redirects Chimeric Antigen Receptor T Cells for Specific and Effective Targeting of Glioblastoma

**DOI:** 10.1101/2020.01.24.918888

**Authors:** Dongrui Wang, Renate Starr, Wen-Chung Chang, Brenda Aguilar, Darya Alizadeh, Sarah L. Wright, Xin Yang, Alfonso Brito, Aniee Sarkissian, Julie R. Ostberg, Yanhong Shi, Margarita Gutova, Karen Aboody, Behnam Badie, Stephen J. Forman, Michael E. Barish, Christine E. Brown

## Abstract

While chimeric antigen receptor (CAR) T cells have demonstrated antitumor activity against glioblastoma (GBM), tumor heterogeneity remains a critical challenge. To more effectively target heterogeneous GBMs, we report the development of a novel peptide-based CAR exploiting the GBM-binding potential of chlorotoxin (CLTX). CLTX bound a greater proportion of tumor cells than GBM-associated antigens EGFR, HER2 and IL13Rα2. CAR T cells bearing CLTX as the targeting domain (CLTX-CAR), mediated potent *in vitro* and *in vivo* anti-GBM activity, and efficiently targeted tumors lacking expression of other GBM-associated antigens. Importantly, CLTX-CAR T cells exhibited no observable off-target effector activity against normal cells, or when adoptively transferred into mice. Effective targeting by CLTX-CAR T cells required cell surface expression of matrix metalloproteinase-2 (MMP-2). Our results are the first demonstration of a peptide toxin utilized as a CAR targeting domain, expanding the repertoire of tumor-selective CAR T cells with the potential to reduce antigen escape.

**One Sentence Summary:** Chimeric antigen receptors incorporating chlorotoxin as the tumor targeting domain recognize and kill glioblastoma with high specificity and potency.

## Introduction

Glioblastoma (GBM) is the most common type of primary brain tumor. Despite the increasingly aggressive treatments incorporating surgery, chemotherapy and radiotherapy, the survival of GBM patients has been only modestly improved over the last several decades (*1, 2*). The poor prognosis for patients with GBMs has prompted the development of advanced therapies, among which is immunotherapy using T cells engineered with chimeric antigen receptors (CARs) (*3, 4*). CAR T cell therapy redirects the cytotoxic activity of T lymphocytes independent of MHC restriction and without need for antigen priming. This cellular therapy, therefore, provides a strategy to generate *de novo* antitumor immunity, which may overcome the challenges of low mutational burden and lack of immunogenicity in tumors such as GBMs with low mutational burdens (*5, 6*). We and others have demonstrated that CAR T cell therapy can be successfully translated for the treatment of GBM (*7–10*), demonstrating safety, evidence for antitumor activity and in one case the potential for mediating complete tumor remission (*8*).

Despite encouraging evidence of clinical safety and bioactivity for GBM-targeted CAR T cells, the overall response rates have been unsatisfyingly low, especially as compared to the remarkable clinical responses observed against B cell malignancies (*11, 12*). One of the major obstacles complicating CAR T cell therapeutic efficacy is tumor heterogeneity, which is particularly significant in GBMs. The classification of GBM subtypes has illustrated the heterogeneity across patients, and more recent studies using single cell sequencing also revealed considerable genetic variations among intratumoral subpopulations (*13, 14*). Efforts to develop CAR T cells for GBMs, therefore, must consider this high degree of heterogeneity. We have demonstrated that IL13 receptor α2 (IL13Rα2) expression is associated with a mesenchymal phenotype and poorer prognosis of GBM patients (*15*). For IL13Ra2-positive tumors, its expression has also been verified on tumor-initiating glioma stem-like cells, and expression is associated with tumor invasiveness (*16*). However, after treating with IL13Rα2-targeted CAR T cells, instances of tumor recurrence with loss and/or reduced expression of IL13Rα2 has been observed (*8, 17*). Similar results were also reported following EGFR variant III (EGFRvIII)-targeted immunotherapies, with the down-regulation of EGFRvIII expression in the tumors harvested post-therapy (*10, 18*). Indeed, GBMs are able to rapidly adapt to therapies, resulting in relapse with distinct intratumoral cellular profiles (*19*). Given the promising clinical results of CAR therapy against B cell malignancies by targeting CD19 which is broadly expressed by all B cell lineages (*20–22*), the development of more effective GBM-targeting CAR designs is expected to benefit from improving broad tumor recognition. To date, this has remained elusive due to the scarcity of antigen candidates that are both widely expressed and highly tumor-specific, especially given the extreme risk of off-tumor toxicities due to the critical location of these tumors within the brain.

An opportunity to extend the repertoire of target antigens amenable to CAR T cell therapy is presented by the tumor-binding potential of some naturally-derived molecules (*23*). One example is chlorotoxin (CLTX), a 36-amino acid peptide isolated from the venom of the death stalker scorpion (*24*). The GBM-binding potential of CLTX was first identified through conjugation with the radioisotope ^131^I (*25*). While the precise cell surface receptor for CLTX on GBM cells remains unclear, CLTX binding impairs GBM cell migration and invasiveness (*26, 27*). Importantly, despite the tumor-binding potential, CLTX itself elicits minor toxic reactions against both tumor and normal tissues (*28, 29*). Therefore, one line of research has been focused on incorporating other cytotoxic agents with CLTX, aiming for tumor-specific delivery. An early clinical study using ^131^I-labeled CLTX successfully mediated radiotherapy against post-resection residual tumor without major neurotoxicity (*28*). CLTX has also been used to coat a variety of delivery vehicles to administer chemotherapy drugs as well as small interfering RNAs (siRNAs) (*30, 31*). More recently, the potential of CLTX to deliver fluorescent signal specifically to tumors (named “Tumor Paint”) has been demonstrated in preclinical models and is currently under clinical investigation (*29, 32, 33*). These studies have indicated that CLTX can be manipulated and harnessed for specific tumor targeting.

Here we report the preclinical development of a novel peptide-based CAR using CLTX as the tumor-targeting domain. CLTX specifically bound a panel of primary GBM cells, with a more inclusive pattern than expression of other GBM-associated antigens. We show that CLTX-CAR T cells incorporating an IgG4(EQ) spacer and a CD28 co-stimulatory signal, were active against GBMs with distinct phenotypes, and elicited antitumor immune responses without major off-tumor toxicities in mouse models with human GBM xenografts. Therefore, our results suggest that CLTX-CAR T cells have the potential to effectively mediate GBM eradication, and represent an advanced approach to CAR design for broad and specific tumor targeting.

## Results

### CLTX binds to a broad variety of GBM cells

Previous studies have documented the GBM-selective binding properties of CLTX (*25, 34*). However, CTLX binding, in relation to either other tumor associated antigens or to GBM subpopulations, has not been examined. We first evaluated CLTX binding to freshly-dissociated tumor cells from surgical resection specimens. These primary brain tumor (PBT) cells were examined by flow cytometry for binding of Cy5.5-conjugated CLTX peptide (CLTX-Cy5.5) and compared with expression of IL13Rα2, HER2 and EGFR, three targets being clinically evaluated in CAR T cell therapies for GBMs (*10, 17, 35*). Strong CLTX-Cy5.5 fluorescence was observed for almost all patient tumors, with greater than 80% of cells binding CLTX (Fig. 1A and 1B, left panel). Across 22 tumor samples from 15 different patients, only two (PBT114 and PBT131) showed CLTX-Cy5.5 binding in less than 30% of total cells. At the same time, expression of immunotherapy targets IL13Rα2, HER2 and EGFR varied widely between patient tumors. CLTX-Cy5.5 binding appeared independent of other antigens, and was observed on tumors with both high and low expression of IL13Rα2, HER2 and EGFR (See representative flow cytometry plots in Fig. 1A). We also examined CLTX-Cy5.5 binding to low-passage patient-derived GBM tumor sphere (PBT-TS) lines expanded under conditions favoring a cancer stem cell-like phenotype (*36–38*). Similar to dissociated primary GBM cells, 18 out of 19 PBT-TS lines showed greater than 80% CLTX-Cy5.5 binding (Fig. 1B, right panel and Fig. S1A), including the TS lines which displayed negligible expression of IL13Rα2, HER2 and EGFR (PBT003-4-TS, PBT009-TS). To evaluate CLTX binding in engrafted tumors, GBM orthotopic xenografts were tested by fluorescent microscopy using a biotin-conjugated CLTX peptide. Consistent with the analyses of freshly resected patient tumors, CLTX displayed consistent binding to five of five engrafted PBT-TS GBM tumors, and marked a greater proportion of tumor cells as compared to the expression of IL13Rα2 and EGFR (Fig. 1C and Fig. S1B). Taken together, these studies confirmed the capacity of CLTX to bind to a high percentage of patient GBM tumors (20 of 22 freshly-dissociated GBM samples), as well as to the majority of GBM cells within each tumor.

**Fig. 1.**
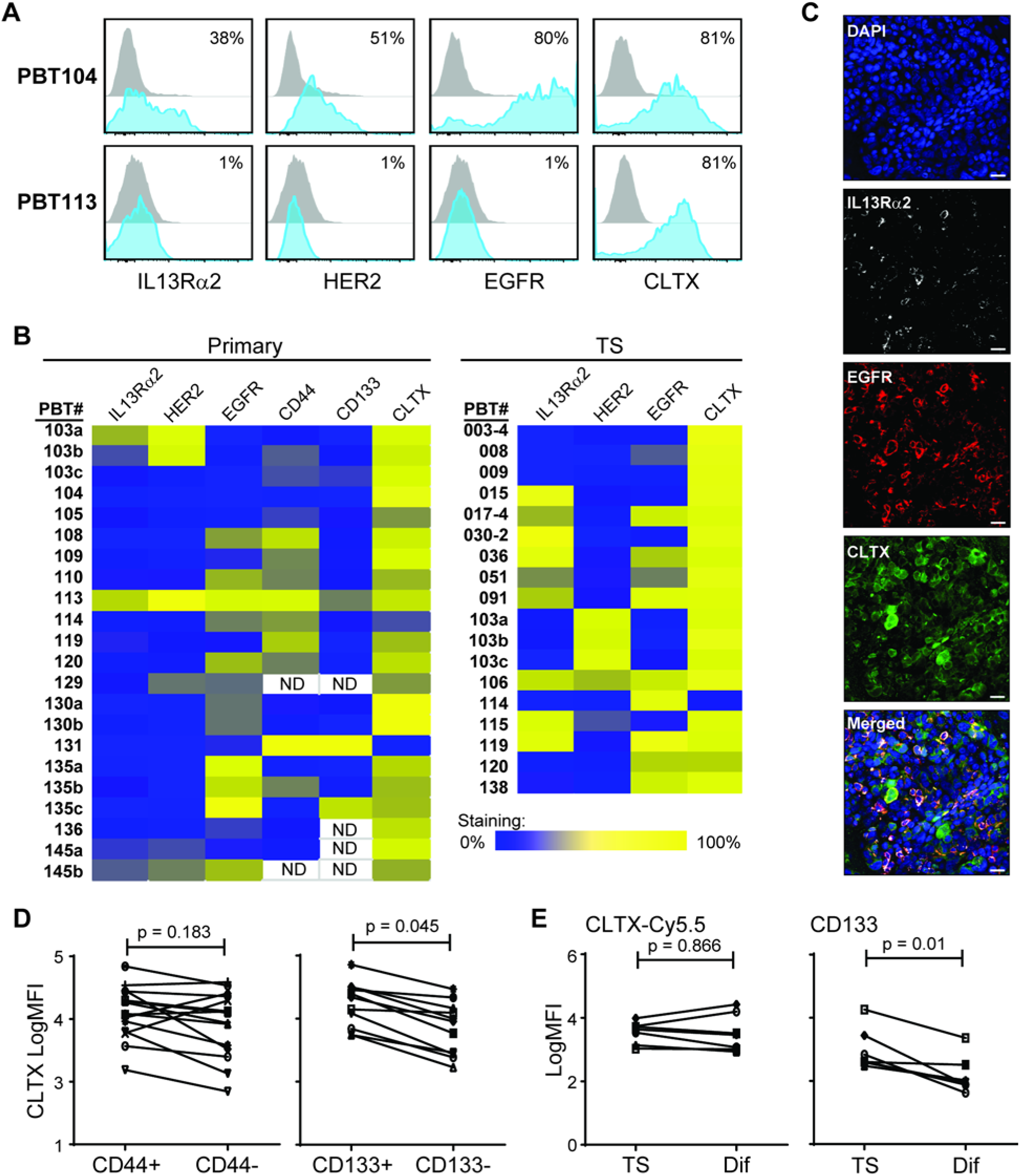
CLTX binds broadly to primary GBM samples. (**A**) Freshly dispersed viable patient brain tumor (PBT) GBM cells (gated as DAPI-, CD45-, CD31-) were immunostained for expression of IL13Rα2, HER2, EGFR, or binding by Cy5.5-conjugated CLTX. Percentages of stained cells (blue) compared to control staining (grey) are indicated in each histogram. (**B**) Summary of staining results (% positive as in (A)) for 22 primary PBTs (*left*) or 18 PBT tumor spheres (TS) (*right*). ND, not done. (**C**) Representative phenotype of a GBM xenograft established by stereotactic injection of 1 x 10^5^ PBT106 TS cells into the right forebrain of an NSG mouse. Tumor-bearing mouse brain was harvested 97 days after cell injection, and paraffin sections were stained with antibodies against IL13Rα2, EGFR, and biotin-conjugated CLTX. Staining was visualized by fluorochrome-conjugated secondary antibody staining and DAPI to identify nuclei. Scale bars: 20μM. (**D**) Freshly-dispersed GBM samples separated into CD44+ and CD44- (*left*) or CD133+ and CD133- (*right*) subpopulations were examined for differences in CLTX-Cy5.5 staining, measured as mean fluorescence intensities (MFI). Only primary PBT samples with >20% of CD44+ or CD133+ fractions were analyzed. (**E**) PBT-TS lines, maintained in neural stem cell medium (TS) or differentiation medium (Dif) for 14 days, were evaluated for CLTX-Cy5.5 and CD133 staining (MFI). (**D, E**) p values are indicated using paired two-way Student’s t-test.

Cells within GBM tumors are highly heterogeneous, composed of phenotypically and functionally distinct subpopulations. Within GBM tumors, stem-like cells (GSCs) display self-renewal and tumor-initiation capacity (*39*), identifying this population as a therapeutically important component of a GBM-targeted CAR T cell therapy (*16, 35, 40*). Hence, we examined CLTX binding with respect to this stem cell-like population. First in freshly-dissociated primary PBTs, we distinguished GSCs from other GBM cells by expression of surface markers CD133 or CD44 (*41, 42*). In the aggregate, we observed that CLTX-Cy5.5 binding was somewhat higher in CD133+ GSCs, but also evident in more differentiated CD133-cells, whereas no significant difference was observed between CD44+ and CD44-GBM cells (Fig. 1D). In another approach, we used PBT-TS lines and varied culture conditions to favor GSCs or alternatively to promote differentiation (*16*). Differentiation led to reduced expression of GSC-marker CD133; however, CLTX-Cy5.5 binding was not affected in comparison to GSCs (Fig. 1E). These studies demonstrated that CLTX binding, while showing some preference for CD133+ GSCs in freshly dispersed tumor samples, remains robust on both stem-like and more differentiated GBM cells. Together, these studies demonstrated the broad GBM binding potential of CLTX, providing rationales for investigating its use to redirect CAR T cell immunotherapy.

### CLTX-CAR T cell potency can be optimized through modifying non-targeting domains

We next sought to design a CAR incorporating the CLTX peptide as the tumor targeting domain. The initial CLTX-CAR was generated using the backbone of our CD19-targeted CAR that has shown safety and clinical activity against B-cell malignancies (*43*). This CLTX-CAR construct is comprised of the CLTX peptide, an IgG4Fc(EQ) spacer, and a CD28 costimulatory domain (Fig. 2A), and is referred to as CLTX-EQ-28ζ. Cytotoxicity of CLTX-EQ-28ζ CAR T cells was evaluated against a panel of GBM PBT-TS lines. CLTX-EQ-28ζ CAR T cells conjugated with tumor cells within 2 hours and formed immunological synapse-like structures (Fig. 2B). Co-culture with GBM cells stimulated CLTX-EQ-28ζ CAR T cells to up-regulate T cell activation markers CD69 and 4-1BB (CD137), and to degranulate as measured by cell-surface CD107a expression (Fig. 2C and Fig. S2A). Further, CLTX-EQ-28ζ CAR T cells efficiently killed GBM cells in 48-hour co-culture assays at an effector to target (E:T) ratio of 1:4 (Fig. 2D). Of note, CLTX-EQ-28ζ CAR T cells were effective against four unrelated PBT-TS lines (Fig. 2D) that exhibited distinct IL13Rα2, HER2 and EGFR expression profiles (Fig. 1B and Fig. S1A). In particular, while IL13Rα2-targeted CAR T cells failed to respond to the PBT-TS lines with low/negative IL13Rα2 expression (PBT003-4-TS, PBT138-TS), CLTX-EQ-28ζ CAR T cells recognized and responded to all four TS lines (Fig. 2, B-D). Further, focusing on PBT003-4-TS, which showed negligible expression of IL13Rα2, HER2 or EGFR (Fig. S1A), we demonstrated that co-culture with CLTX-EQ-28ζ, but not the combination of CAR T cells targeting all three other antigens, mediated tumor cell elimination (Video.S1 and S2). Lentiviral transduction of PBT003-4-TS to over-express IL13Rα2, HER2 or EGFR did not reduce CLTX-EQ-28ζ cytotoxic activity (Fig. S2, B and C). We thus concluded that CLTX-EQ-28ζ was able to recognize and target GBM cells, and its cytotoxicity is consistent with the broad GBM-binding potential of the CLTX peptide while independent of other GBM-associated antigens.

**Fig. 2.**
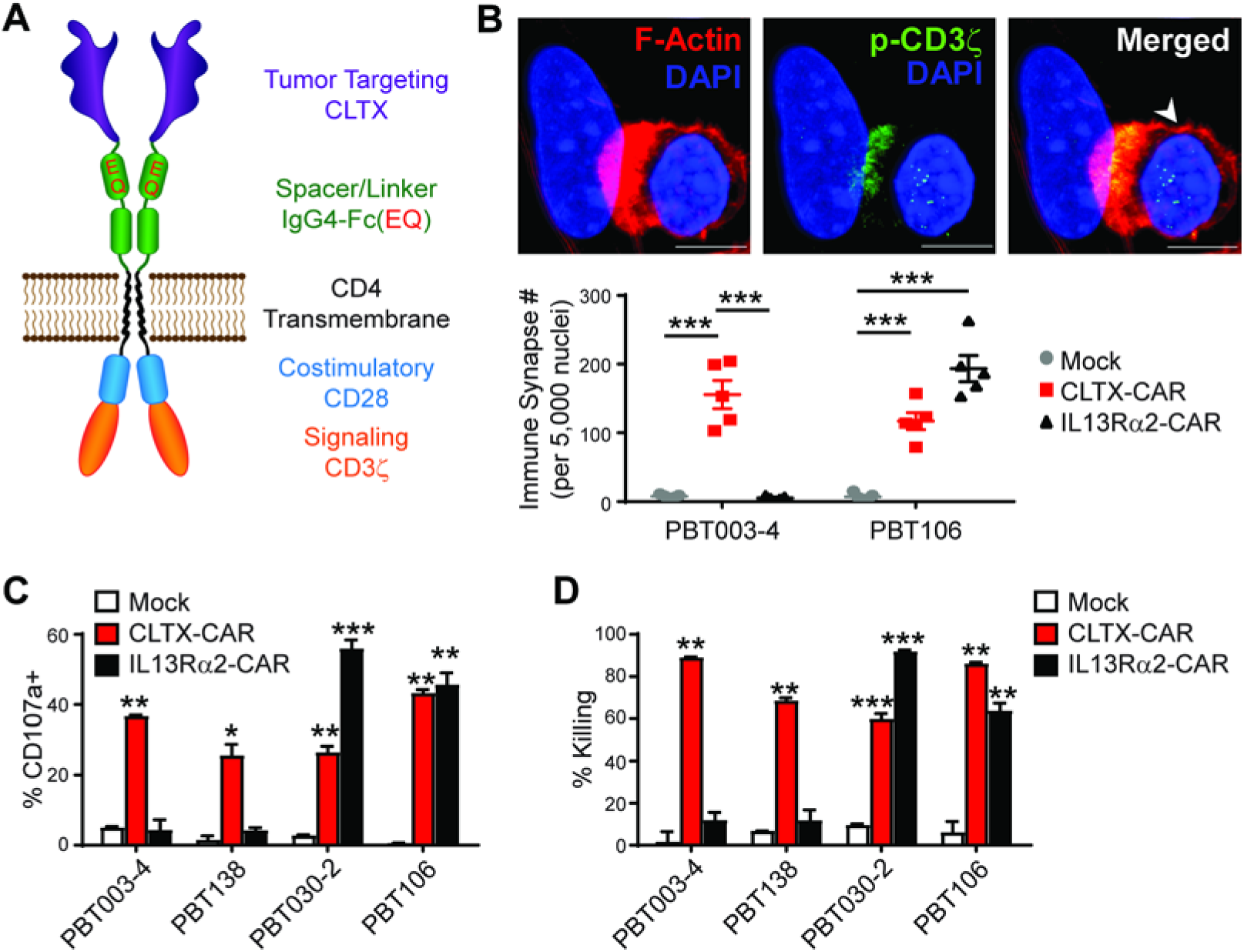
Effector activity of CLTX-CAR T cells. (**A**) Diagram of a CAR containing a tumor targeting CLTX domain, an IgG4-Fc spacer domain with EQ mutations, a CD4 transmembrane domain, and intracellular signaling domains (CD28 and CD3ζ). (**B**) Immunological synapse formation at 2h after adding CLTX-CAR T cells to GBM cells dissociated from PBT003-4-TS (IL13Rα2-) or PBT106-TS (IL13Rα2+) (E:T=1:1). *Top*, a representative image of immunological synapse indicated by co-localization of phosphorylated-CD3ζ (p-CD3ζ) and polarized F-actin accumulation at the interface between the CLTX-EQ-28ζ CAR T cell and the tumor cell. Arrowhead: T cell; scale bars: 5 μM. *Bottom*, Number of immunological-synapses per 5,000 tumor nuclei, when mock-transduced T cells, CLTX-CAR T cells, or IL13Rα2-CAR T cells, were co-cultured with different GBM cells. Five different fields were counted for each group. Mean±SEM is plotted; ***, p<0.001. (**C**) Degranulation of CLTX-CAR T cells (% CD107a) after co-culture with GBM cells from different PBT-TS lines. Mock-transduced T cells, CLTX-CAR T cells or IL13Rα2-CAR T cells were stimulated with either IL13Rα2-negative (PBT003-4, PBT138) or IL13Rα2+ (PBT030-2, PBT106) GBM cells at a 1:1 E:T ratio for 5h, and CD3/CD19t+ gated cells were analyzed for surface CD107a expression as a marker of degranulation. Mean ± S.E.M of % CD107a+ cells in duplicate wells are depicted. (**D**) GBM cells were co-cultured with mock T cells, CLTX-CAR T cells or IL13Rα2-CAR T cells at an effector:target (E:T) ratio of 1:4 for 48 h. CAR T cell activity is shown by the percentage of target cells eliminated. (**C, D**) *, *p*<0.05; **, *p*<0.01; ***, *p*<0.001 compared with mock T cells using one-way ANOVA with Bonferroni’s Multiple Comparison Tests. All PBT numbers indicate PBT-TS lines.

The cytolytic activity of CAR T cells is greatly influenced by the design of regions outside of the antigen-targeting domain, including the spacer (*44*) and the co-stimulatory domains (*45–47*). Using the CLTX-EQ-28ζ CAR as a reference, we first addressed the impact of spacer length by generating CAR constructs to compare IgG4Fc(EQ) (239 amino acids) with three shorter spacers: IgG4-Fc with the CH2-domain deleted (ΔCH2) (129 amino acids), CD8 hinge (CD8h) (44 amino acids), and a short synthetic linker (L) (10 amino acids) (Fig. 3A). We observed that CAR T cell-mediated tumor killing was greatly reduced with ΔCH2 or L spacers, while the CD8h spacer retained CAR function similar to that of IgG4Fc(EQ) (Fig. 3B). We next evaluated the contributions of co-stimulatory signals by generating CARs bearing CD28 or 4-1BB co-stimulatory domains in the contexts of spacers IgG4Fc(EQ) and CD8h (Fig. 3A). Intriguingly, CAR T cells incorporating a CD28 co-stimulation domain consistently showed higher T cell effector function as compared with CARs incorporating a 4-1BB co-stimulation domain (Fig. 3C). We then evaluated the mechanism underlying the functional differences across CLTX-CAR constructs. Notably, for the six CLTX-CAR constructs evaluated, we observed consistent tumor-dependent activation across multiple assays including killing potency (% killing), extent of degranulation (% CD107a+), and secretion of cytokine IFNγ (Fig.3, B and C, and Fig. S3A). Moreover, consistent with the potent cytotoxic effector function of CLTX-EQ-28ζ and CLTX-CD8h-28ζ CARs, both CAR T cells also showed expression of T cell activation markers 4-1BB and CD69 (Fig. S3B), and inhibitory molecule PD-1 which is also linked to T cell activation (Fig. S3C). These results therefore suggested that differences in initial activation were the major contributor to variations in effector function across various CLTX-CAR designs.

**Fig. 3.**
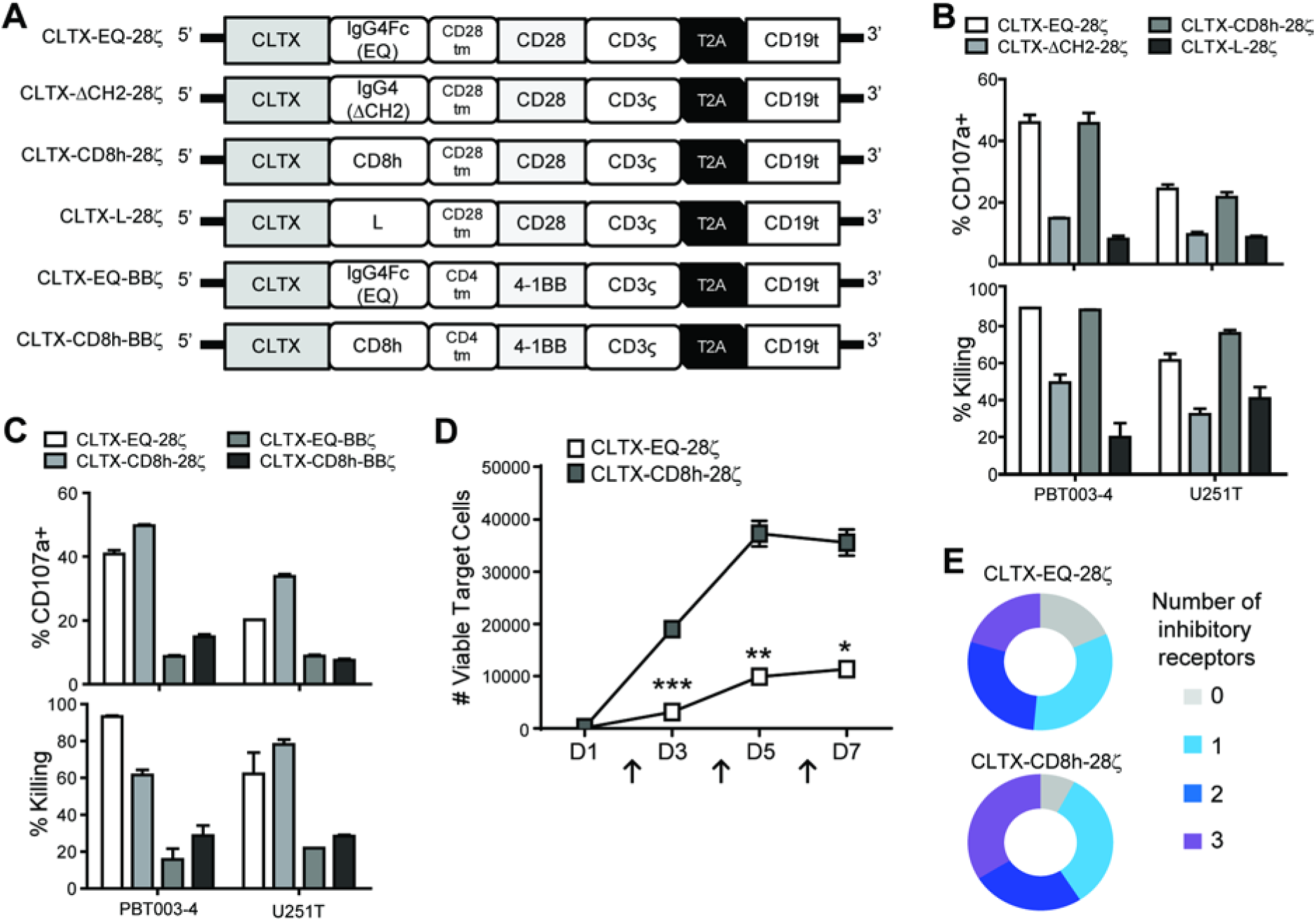
Optimization of CLTX-CAR design for effector potency. (**A**) Schemas of CLTX-CARs bearing different spacers and/or co-stimulatory signals. (**B**) Degranulation (% CD107a+; top) and cytotoxicity (% target cell killing; bottom) for CLTX-CAR T cells with CD28 costimulatory domains but different spacers, tested against PBT003-4-TS or U251T GBM cells. (**C**) Degranulation (top) and cytotoxicity (bottom) of CLTX-CAR T cells with IgG4Fc(EQ) or CD8h spacers, and CD28 or 4-1BB co-stimulatory domains, tested against PBT003-4-TS or U251T GBM cells. (**D**) CLTX-EQ28ζ and CLTX-CD8h28ζ CAR T cells were co-cultured with PBT003-4-TS GBM cells (4,000 CAR+ cells, 16,000 tumor cells), and repetitively challenged with 32,000 tumor cells every 48 hours (arrows, D2, 4 and 6). Numbers of remaining viable tumor cells were quantified at the indicated time points during the rechallenge assay. Mean± SEM of duplicate wells are shown; *, p < 0.05, **, p < 0.01, and ***, p < 0.001 using an unpaired Student’s t-test. (**E**) At Day 4 of rechallenge, CLTX-EQ28ζ and CLTX-CD8h28ζ CAR T cells were analyzed for T cell exhaustion markers and co-expression of PD-1, LAG-3 and TIM-3. All PBT numbers indicate PBT-TS lines.

We next returned to the comparison of effector potency between CLTX-EQ-28ζ and CLTX-CD8h-28ζ CAR T cells. First, screening degranulation and cytokine-production against a panel of 10 PBT-TS lines, we observed that CLTX-EQ-28ζ CAR T cells displayed higher degranulation, but lower IFNγ production as compared to CLTX-CD8h-28ζ CAR T cells (Fig.S3, D and E). Next, the capacities of CLTX-EQ-28ζ and CLTX-CD8h28ζ CAR T cells to maintain long-term antitumor activity were evaluated in a repetitive tumor challenge assay (*48*). We observed that CLTX-EQ-28ζ, but not CLTX-CD8h-28ζ, CAR T cells retained activity through multiple rounds of tumor challenge (Fig. 3D). Since both CLTX-EQ-28ζ and CLTX-CD8h-28ζ CAR T cells significantly upregulated PD-1 expression upon tumor stimulation (Fig. S3C), we then assessed co-expression of three T cell inhibitory receptors (PD-1, LAG-3 and TIM-3) which together marks exhausted T cells (*48, 49*), and observed that the absence of durable effector function in CLTX-CD8h-28ζ CAR T cells was associated with an exhausted phenotype featuring co-expression of multiple inhibitory receptors (Fig. 3E). Considering all of these observations about effector potency, we adopted the CLTX-EQ-28ζ CAR as the optimal design for subsequent evaluation.

### CLTX-CAR T cells mediate antitumor activity against established GBM xenografts

To test the *in vivo* antitumor activity of CLTX-EQ-28ζ CAR T cells, xenograft tumors were established using two patient-derived PBT-TS lines with distinct antigen expression patterns: PBT003-4-TS and PBT106-TS (Fig. S1A). Against subcutaneously engrafted GBMs, intratumoral administration of CLTX-EQ-28ζ CAR T cells resulted in tumor regression, while tumors injected with mock-transduced T cells displayed growth kinetics similar to tumor-only controls (Fig. S4A). Next, the antitumor function of CLTX-EQ-28ζ CAR T cells was evaluated in orthotopic GBM models in which PBT-TS lines were stereotactically implanted into the brains of immunodeficient mice. After engraftment, the brains were subjected to intracranial injection of CAR T cells (*50*). Consistently with the subcutaneous tumor model, intracranially-administered CLTX-EQ-28ζ CAR T cells, but not mock T cells, controlled tumor growth and prolonged the survival of mice bearing PBT003-4-TS or PBT106-TS tumors (Fig. 4, A-D).

**Fig. 4.**
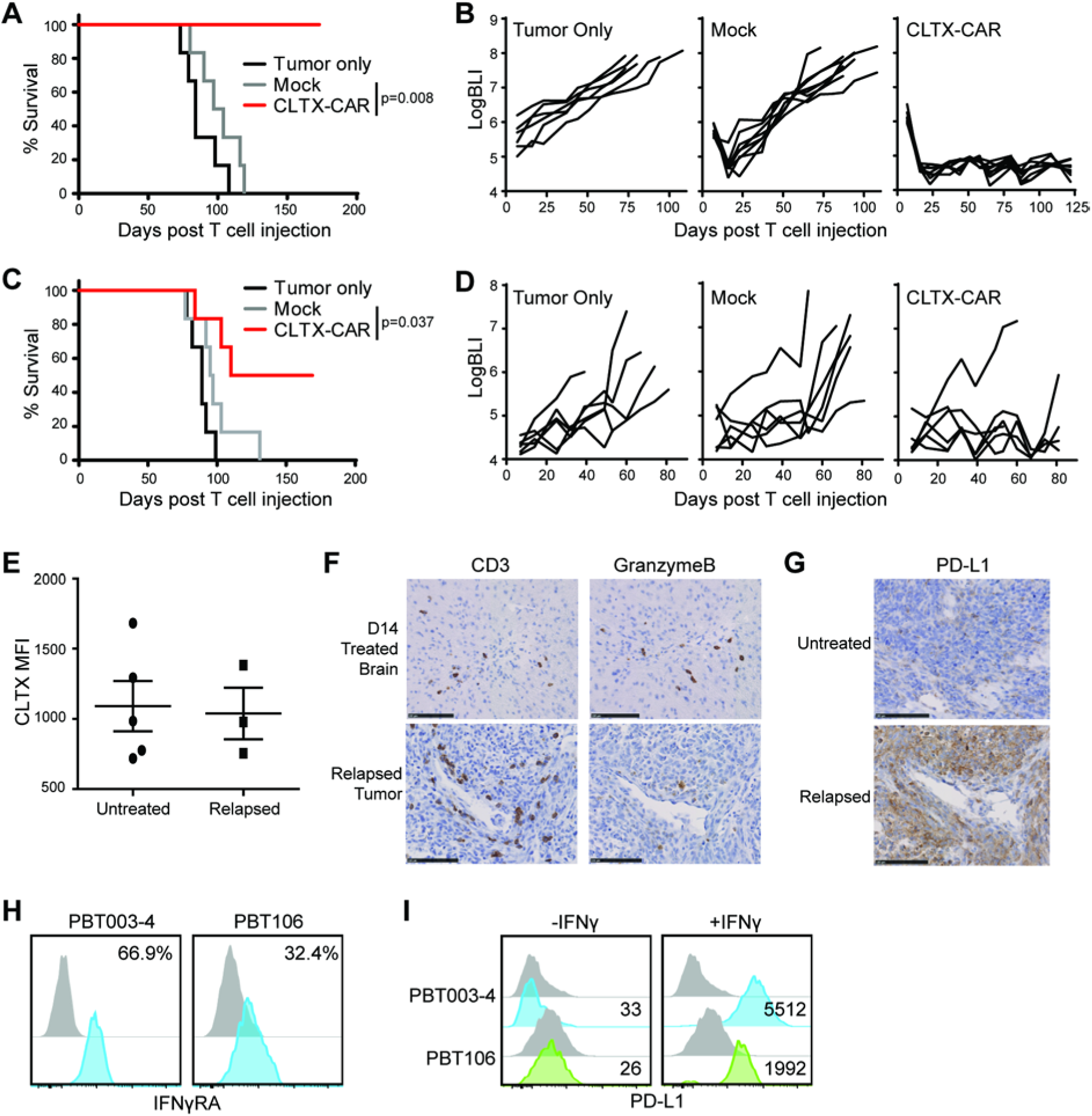
*In vivo* antitumor activity of CLTX-CAR T cells. ffLuc^+^ PBT106-TS (**A**,**B**) or PBT003-4-TS (**C**,**D**) GBM cells were stereotactically implanted into the right forebrain of NSG mice (1 × 10^5^ cells/mouse). On day 8 post tumor implantation, mice (n=6-7 per group) received either no treatment (Tumor only); intracranial treatment with 1×10^6^ mock-transduced T cells (Mock) or CLTX-EQ-28ζ CAR+ T cells (CLTX-CAR). (**A**,**C**) Kaplan Meier survival analysis with Log-rank (Mantel Cox) test comparing CLTX-EQ28ζ CAR T cell and Mock T cell treated groups. (**B**,**D**) Tumor volumes over time monitored using bioluminescent imaging. (**E**) Tumors harvested from NSG mice bearing PBT003-4-TS GBM xenografts that were either untreated (n = 5) or relapsed from those treated with CLTX-CAR T cells (n = 3), were dissociated into single cell suspensions and stained with CLTX-Cy5.5. Lines indicate mean MFI ± SEM. (**F**) Immunochemical staining for CD3 and granzyme B on mouse brain sections at D14 post T cell injection and 7 days after tumor clearance (*top*), or on the relapsed tumor (*bottom*). (**G**) PD-L1 staining on untreated (*top*) or relapsed (*bottom*) PBT003-4-TS tumors. (**H**) Expression of IFNγ Receptor A (IFNγRA) on GBM cells dissociated from PBT003-4-TS and PBT106-TS. Percentages of immunoreactive cells (blue) above that of isotype control staining (grey) are indicated in each histogram. (**I**) PD-L1 surface expression on PBT003-4-TS and PBT106-TS GBM cells with and without IFNγ treatment. Numbers indicate PD-L1 MFI above isotype control. All PBT numbers indicate TS lines.

Of note, the responses of engrafted PBT003-4-TS and PBT106-TS tumors to CLTX-CAR T cell therapy were not equivalent. After CAR T cell administration, all mice bearing PBT106-TS tumors remained tumor-free for over 170 days, while only a subset of PBT003-4-TS tumor-bearing mice achieved similar long-term tumor eradication (Fig. 4A-D and Fig. S4B). To investigate the etiology of this variable tumor response against different GBM models, we examined recurrent PBT003-4-TS tumors post CLTX-EQ-28ζ therapy. We first observed that CLTX-Cy5.5 binding on these recurrent tumors was similar to untreated tumors (Fig. 4E), suggesting that antigen escape did not account for tumor relapse. Investigating the behavior of antitumor T cells within these orthotopic tumors, we found that during primary antitumor response, granzyme B-expressing T cells were detected 14 days following adoptive transfer (Fig. 4F top row). By comparison, while CLTX-EQ-28ζ T cells still persisted in the recurrent tumors, these T cells expressed only low levels of granzyme B (Fig. 4F bottom row). At the same time, these relapsed PBT003-4-TS tumors displayed increased expression of PD-L1 as compared to untreated tumors (Fig. 4G). PD-L1 induction on tumors post CAR treatment was consistent with a high IFN-γ receptor A (IFNγRA) expression in GBM cells from PBT003-4-TS before implantation (Fig. 4H). IFN-γ had been shown to promote PD-L1 transcription through its receptors in other tumors (*51*), and IFN-γ treatment rapidly induced PD-L1 expression by PBT003-4-TS cells (Fig. 4I). Notably, cells from PBT106-TS exhibited lower levels of IFNγRA expression and less robust PD-L1 induction by IFN-γ treatment, as compared to PBT003-4-TS (Fig. 4, H and I). These observations suggested that for some GBMs, adaptive response to immunostimulatory cytokines in tumors and subsequent induction of inhibitory checkpoint receptors, rather than antigen escape, was the main reason for insufficient tumor eradication by CLTX-CAR T cells.

### CLTX-CAR T cells have minimal off-target effects

Prior clinical studies using CLTX to deliver radiation and imaging reagent to tumor sites (NCT00040573, NCT02234297) have not reported significant adverse events, and multiple preclinical studies have shown no off-target toxicity (*29, 32*). Consistent with these reports, we observed limited to undetectable CLTX-Cy5.5 binding to a panel of non-tumor cells, including periphery blood mononuclear cells (PBMCs), human embryonic kidney 293T cells, as well as induced pluripotent stem cell-differentiated astrocytes (iPSC-Ast), neural progenitor cells (iPSC-NPCs) and immortalized fetal brain derived neural stem cell (FB-NSC) line LM-NSC008 (*52–54*) (Fig. 5A). We next concerned that even a low-level of binding could trigger CLTX-CAR T cell activation. Therefore, we evaluated the activation potential of CLTX-EQ-28ζ CAR T cells during co-culture with these normal cells. As indicated by degranulation and *in vitro* cytotoxicy (Fig. 5, B and C), CLTX-EQ-28ζ CAR T cells did not target PBMC, 293T, iPSC-Ast, iPSC-NPC and FB-NSC, exhibiting only background levels of effector activity equivalent to mock transduced T cells. Further, the observation that CLTX-Cy5.5 binding is conserved between mouse and human GBM cells (see Fig. 5D for examples of CLTX-Cy5.5 binding to mouse KR158 and GL261 GBM cells) supports the relevance of murine models for assessing CLTX-CAR T cell off-tumor toxicities. In mouse brains bearing GBM xenograft tumors, CLTX-Cy5.5 bound to tumor cells, but not normal brain tissue, delineating the xenograft-tumor border (Fig. 5E). In addition, intratumoral administration of CLTX-EQ-28ζ CAR T cells in GBM-bearing mice showed no evident toxicity, as indicated by close examination of multiple organs including brain, kidney, liver, spleen, intestine and colon (Fig. S5A). The potential for systemic toxicity was further assessed by intravenous injection of 50×10^6^ CLTX-EQ-28ζ CAR T cells into normal NSG mice. At this high dose, CLTX-CAR T cells were well-tolerated; all animals remained alert and active, and showed no changes in body weight (Fig. 5F). Taken together, these observations strongly support that CLTX-EQ-28ζ CAR T cells specifically target GBM cells while sparing normal tissues.

**Fig. 5.**
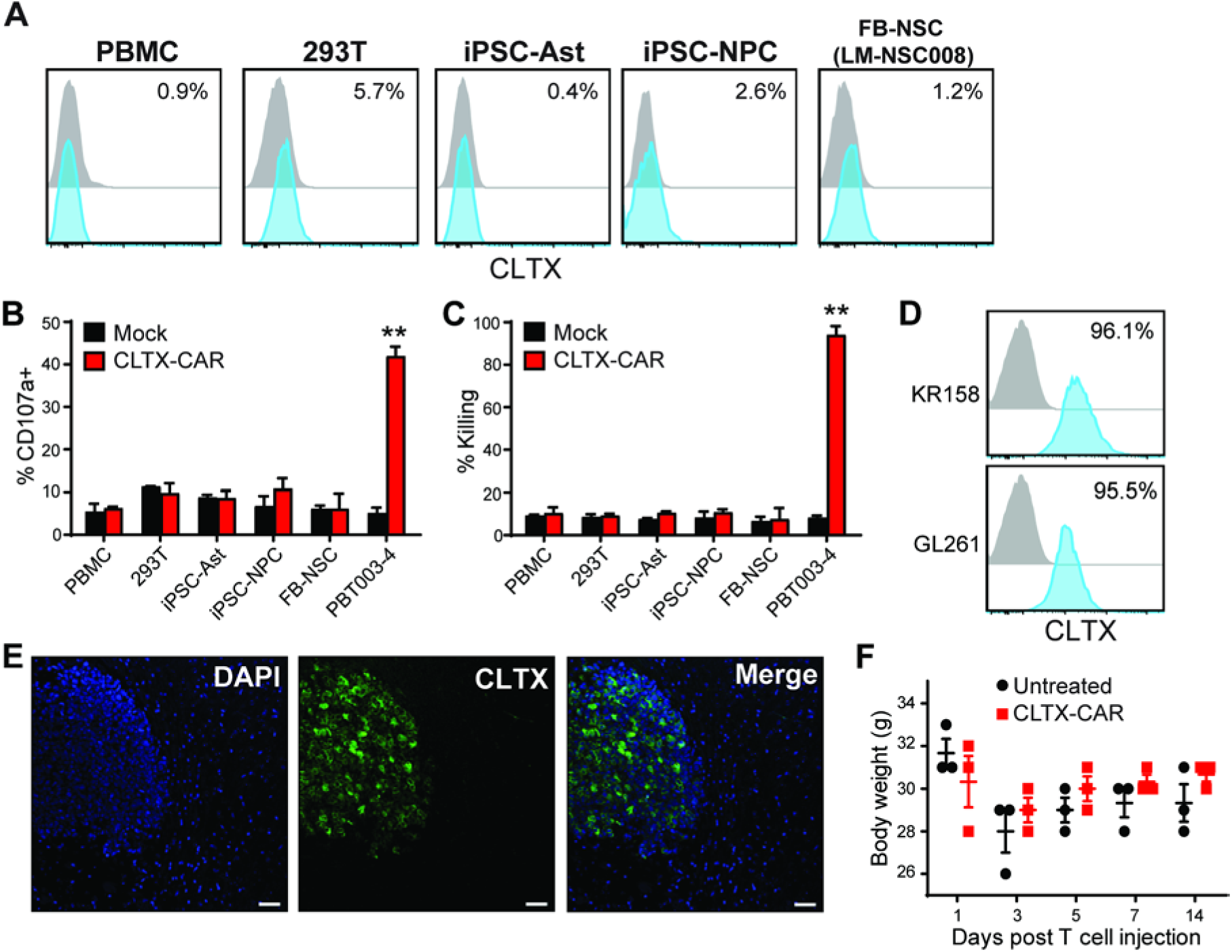
Off-target evaluation of CLTX-CAR T cells. (**A**) CLTX-Cy5.5 staining on viable peripheral blood mononuclear cells (PBMC), human embryonic kidney 293T cells, iPSC-derived astrocytes (iPSC-Ast), neural progenitor cells (iPSC-NPC) or human fetal brain neural stem cells (FB-NSC line LM-NSC008) evaluated by flow cytometry. Percentages of immunoreactive cells (blue) above control (grey) are indicated in each histogram. (**B-C**) Degranulation (**B**) and cytotoxicity (**C**) of mock-transduced (Mock) or CLTX-EQ-28ζ CAR+ (CLTX-CAR) T cells against non-malignant cell lines, with PBT003-4-TS GBM cells as a positive control. Mean + SEM of duplicate wells are depicted; **, p<0.01when compared with mock-transduced T cells using an unpaired Student’s t-test. (**D**) CLTX-Cy5.5 staining on mouse GBM cell lines. Percentages of positively-stained cells (blue) above that of isotype control (grey) are indicated in each histogram. (**E**) Representative immunofluorescent phenotype of a GBM xenograft established by stereotactically injecting 1×10^5^ PBT106-TS cells into the right forebrain of an NSG mouse. The tumor-bearing mouse brain was harvested 93 days after cell injection, and paraffin sections were stained with DAPI to identify nuclei, and CLTX-Cy5.5 to depict the border between xenograft and normal mouse brain. Scale bars: 20μm. (**F**) Body weights of NSG mice receiving intravenous administration of 5×10^7^ CLTX-EQ-28ζ CAR+ (CLTX-CAR) T cells. Body weights were monitored over 2 weeks and compared with untreated NSG mice. Lines indicate mean ± SEM. All PBT numbers indicate TS lines.

### MMP2-expression is required for CLTX-CAR targeting

CLTX binding has been reported to associate with multiple membrane proteins, including membrane bound matrix metalloproteinase 2 (MMP2), chloride channel CLCN3, and phospholipid protein Annexin A2 (AXNA2) (*26, 27, 55*). With the composition of the CLTX receptor poorly identified, we took advantage of CLTX-CAR T cells to investigate the correlation between the expression of these surface proteins on GBM cells and CLTX-CAR effector activity. Using 15 different PBT-TS lines and iPSC-NPCs (negative control) as CLTX-EQ-28ζ CAR T cell targets, we assessed the correlations of MMP2, CCLN3, and AXNA2 expression with CLTX-CAR T cell activity (degranulation potential). We found that CLTX-EQ-28ζ CAR T cell activity varied between these different GBM target cells (Fig. 6A), and observed a strong correlation between CLTX-CAR T cell degranulation and target cell MMP2 expression (Fig. 6B), but not with expression of CLCN3 or ANXA2 (Fig. S6A). Consistent with this pattern, CLTX-Cy5.5 binding correlated with MMP2 expression on target cells (Fig. 6C), but not with CLCN3 or AXNA2 expression (data not shown). Differences in CLTX-CAR cytotoxic potential against different GBM cells were most evident at an E:T ratio of 1:8 (Fig. S6B), and correlated well with degranulation (Fig. S6C). Importantly, the extent of degranulation did not correlate with IFNγRA (CD119) expression on PBT-TS lines (Fig. S6D), which was previously related to low *in vivo* CLTX-CAR T cell function through induction of PD-L1. These results suggest that for GBM cells, MMP2 is the primary mediator of CLTX-Cy5.5 binding and CLTX-CAR T cell cytotoxicity. To further verify the necessity of MMP2 for CLTX-CAR activation, we knocked down MMP2 in GBM cells using lentiviral shRNA. The knockdown dramatically reduced MMP2 transcript levels and MMP2 secretion, and also resulted in modest decreases in CLCN3 expression and minimal changes in ANXA2 expression (Fig. S6E). When subjected to *in vitro* co-culture, we found that CLTX-EQ-28ζ CAR T cell activation and cytotoxicity were dramatically reduced by shMMP2 transduction (Fig.6, D and E, Fig. S6F). Further, MMP2 knockdown significantly reduced *in vivo* antitumor activity of CLTX-EQ-28ζ CAR T cells against tumors established with both PBT003-4-TS and PBT106-TS (Fig. 6F). Together, these results demonstrate that MMP2 is necessary for CLTX-CAR recognition and activation against tumor targets.

**Fig. 6.**
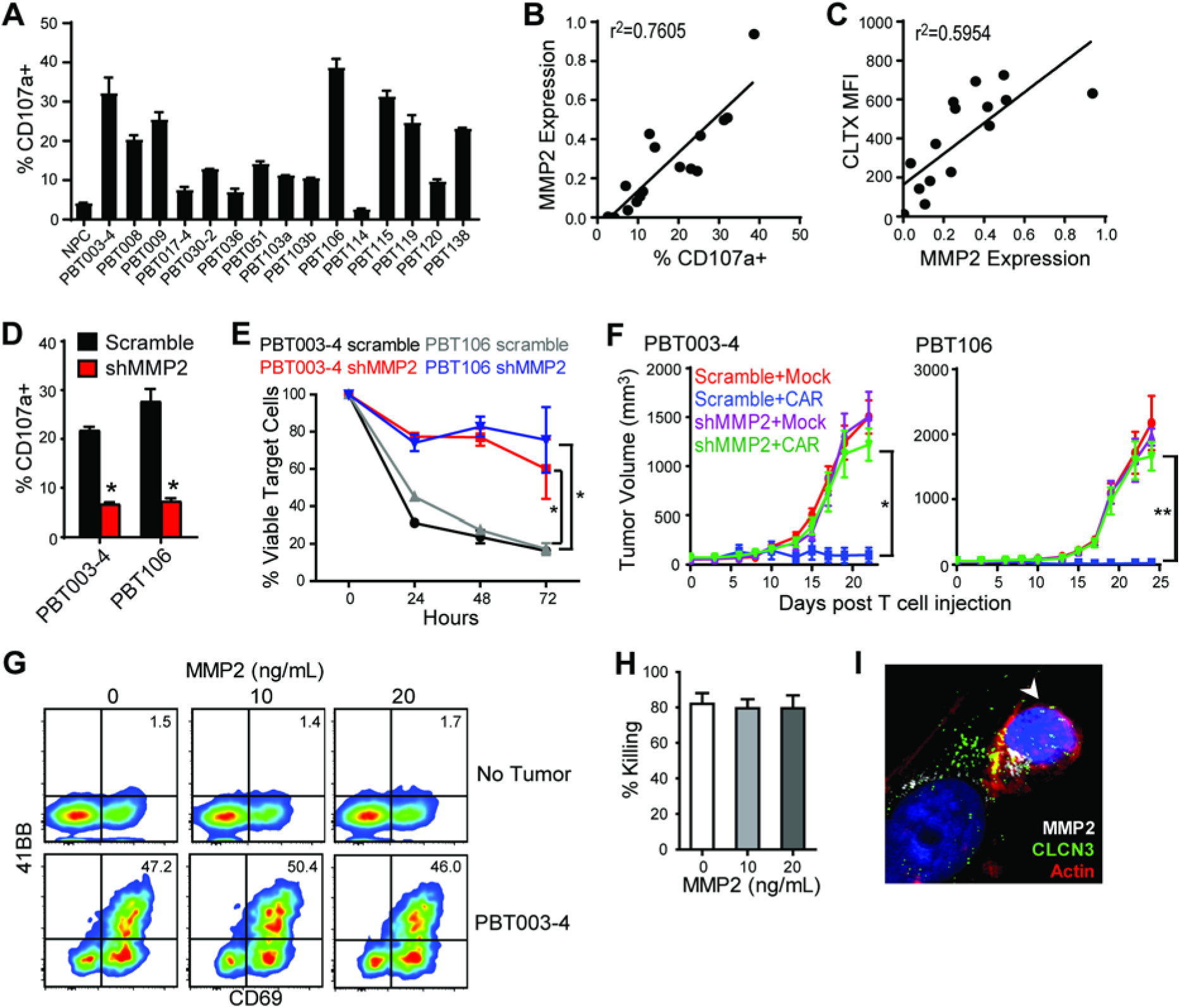
CLTX-CAR T cell effector activity requires MMP2 expression on target cells. (**A**) Degranulation of CLTX-EQ-28ζ CAR+ T cells against iPSC-NPCs and GBM cells from indicated TS lines. Shown are mean ± SEM of duplicate wells. (**B-C**) MMP2 mRNA expression (2^−ΔCt^ compared with ACTIN) in the GBM-TS lines were compared to their corresponding stimulation of CLTX-CAR T cell degranulation (**B**) and CLTX-Cy5.5 staining (**C**); with correlation coefficients (r^2^) indicated in each graph. (**D**) Degranulation of CLTX-CAR T cells tested against PBT003-4-TS or PBT106-TS lines that had been transduced with scramble shRNA or shRNA targeting MMP2 (shMMP2). (**E**) Cytotolytic activity of CLTX-CAR T cells against scramble shRNA or shMMP2 transduced GBM cells over a 72h co-culture (E:T=1:4). (**D, E**) * p < 0.05 when compared with scramble shRNA transduced targets using an unpaired Student’s t-test. (**F**) Scramble shRNA or shMMP2 transduced GBM cells (5×10^6^) were injected subcutaneously into the right flank of NSG mice. Mock-transduced or CLTX-EQ-28ζ CAR+ T cells (3×10^6^) were then injected into the tumors at day 14 (tumor diamater ∼5mm), and tumor size was monitored over time with caliper measurements. * p < 0.05, ** p < 0.01 using one-way ANOVA with Bonferroni’s Multiple Comparison Tests. (**G**) CLTX-CAR T cells were cultured with indicated concentrations of soluble MMP2 in the absence (top row) or presence (bottom row) of PBT003-4 GBM cells (E:T=1:4) for 24h. T cell activation was determined by flow cytometric analysis of 4-1BB and CD69 surface expression. Quadrants were drawn based on control staining, and percentages of double-staining cells are indicated in each histogram. (**H**) Elimination (% killing) of PBT003-4-TS GBM cells by CLTX-CAR T cells after 48 h of co-culture (E:T=1:4) at different concentrations of soluble MMP2. Percentages of tumor killing were calculated using the numbers of viable tumor cells when cultured in the absence of CLTX-CAR T cells; mean ± SEM of duplicate wells are shown. (**I**) Immunofluorescence staining of MMP2, CLCN3 and actin at 2 h after initiating co-culture of CLTX-CAR T cells and PBT003-4 GBM cells (E:T=1:1), arrowhead: T cell. All PBT numbers indicate TS lines.

MMP2 is a secreted matrix metalloproteinase that can either associate with the membrane through interaction with α_v_β_3_ integrin and MMP14 [also known as membrane-type 1 (MT1)-MMP], or exist as a soluble enzyme (*26, 56*). Although our data and others indicate that CLTX binds to the membrane-bound form of MMP2 (*26*), we further investigated whether CLTX-CAR T cells may also respond to soluble MMP2. In the presence of soluble MMP2, we observed no activation of CLTX-CAR T cells (Fig. 6G), and the cytotoxicity of CLTX-CAR T cells against GBM cells was also maintained (Fig. 6, G and H). In addition, we visualized the recruitment of MMP2 to the CLTX CAR T cell-GBM cell immunological synapse (Fig. 6I) but not at the synapse formed by other CAR T cells (Fig. S6G). Meanwhile, we were intrigued to find that CLCN3, although not correlated with CLTX-CAR T cell activation, was also recruited to the immunological synapse (Fig. 6I). Similarly, we observed co-localization of MMP2 and CLCN3 when CLTX peptides were applied to GBM cells (Fig. S6H). These results suggest that binding of CLTX-CAR T cells to a GBM cells depends on expression of MMP2 and involves CLCN3. Recruitment of CLCN3 following MMP2 interaction with CLTX peptide or CLTX-CAR T cells was reminiscent of another study using CLTX-coated liposome (*57*). Together, our results suggested that membrane-bound MMP2 is necessary for CLTX-CAR T cell activation, while soluble MMP2 neither activates nor masks CLTX-CAR function.

## Discussion

This study demonstrates that CLTX, a peptide component of scorpion venom, can be successfully incorporated into a CAR construct to redirect cytotoxic T cells to target GBMs. The antitumor effects of CLTX-CAR T cells were observed in multiple GBM xenograft models. We did not observe off-tumor targeting, and the CLTX-CAR T cells were well tolerated without evident systemic toxicity. Moreover, CLTX-Cy5.5 binding extended across a wider range of freshly-dissociated tumor cells and patient-derived GBM cell lines than expression of antigens currently under consideration as CAR immunotherapy targets (i.e., IL13Rα2, HER2 and EGFR). Our study also provides the first demonstration that a natural toxin peptide can be exploited to generate CAR T cells.

The criteria for selection of GBM-associated immunotherapy targets are particularly stringent because of the sensitivity of the brain to off-target activity or reactions to the therapy (*3*). For the target antigens currently under clinical investigation, one selection criterion is their specific expression on GBM tumors. IL13Rα2 has negligible expression in normal brain and elsewhere (except as a cancer-testis antigen) (*58, 59*). EGFRvIII is the most common mutated form of EGFR restricted to multiple cancers including GBM (*60, 61*). Other antigen targets are over-expressed by tumor GBM cells as compared to normal cells (e.g. EGFR), and HER2, while moderately expressed in some normal tissues, shows only minimal expression on postnatal neurons or glial cells (*62, 63*). Initial development of CLTX-CAR T cells was justified by the demonstrated safety of CLTX itself in previous preclinical and clinical studies (*28*). Specifically, no toxicity was observed in a clinical study using CLTX to deliver ^131^I to tumor sites, or in preclinical studies leading to clinical use of fluorophore-conjugated CLTX as a real-time imaging agent during surgery (*28, 29*). Here we evaluated the potential toxicity of CLTX-CAR T cells in mouse models using multiple strategies, as CLTX binds to mouse glioma cells supporting its relevance for assessing off-tumor toxicities. First, intravenously delivered large-amount of CAR T cells showed no systemic toxicity (Fig. 5F). Further, the regional delivery of CAR T cells into tumor-bearing mice provided a more clinically-relevant condition where the CAR T cells can be activated by GBM tumors to assess off-tumor targeting (Fig. S5). Results of these preclinical studies in mice suggest that CLTX-CAR T cells mediate negligible off-tumor toxicities and provide support for its evaluation in GBM patients.

Tumor recurrence remains a barrier to successful immunotherapy for GBM. Recurrence is commonly associated with the presence of certain cell subpopulation(s), so-called glioma stem cells (GSCs) that are resistant to radiotherapy and chemotherapy, and are characterized by high tumor-initiating potential (*39, 64, 65*). GSCs are usually resistant to conventional therapies, and a critical consideration of GBM-targeted immunotherapy has been the capability to eliminate GSCs (*39, 66*). Indeed, previous studies have demonstrated that GSCs can be as responsive to CAR T cell-mediated cytotoxicity as more differentiated GBM cells (*16, 35, 40*), indicating that GSCs are potentially targetable by CAR T cell therapy despite their intrinsic immunosuppressive properties (*67, 68*). Consistent with this expectation, functional evaluations of CLTX-CAR T cells throughout this study have utilized patient-derived TS lines, which maintain stem-cell like properties under appropriate culture conditions (*69*). On the other hand, we were able to characterize GSCs from freshly-dispersed GBM samples by surface markers CD133 and CD44, representing different GBM molecular subtypes (*65, 70, 71*), and CLTX-Cy5.5 binding appeared to be similar in either GSC or non-GSC populations. All evidence suggests that CLTX-CAR T cells inherit the binding properties of CLTX-Cy5.5 conjugates, and thus will be active against both GSC and non-GSC populations in patient tumors, suggesting the potential to target the “seed” of GBM recurrence.

The choice of co-stimulatory signal has proven to be a critical element of CAR design. CAR T cells with CD28 (*72, 73*) or 4-1BB (*8, 74*), costimulatory signals have been introduced to patients with hematological and solid tumors. In general, compared with CD28, 4-1BB co-stimulation has resulted in slower but more long-lasting T cell activation, as indicated by differential metabolic profiling (*47*), and consistent with CAR T cell expansion dynamics when co-infused into B-ALL patients (*75*). Surprisingly, in this study we found that CLTX-CAR T cells only displayed degranulation and killing antitumor responses when incorporating CD28 co-stimulation, while 4-1BB co-stimulation reduced both short-term and long-term effector activity for CLTX-CAR T cells. One possible explanation may be the nature of CLTX’s interaction with its receptors. Although the binding affinity between CLTX and its receptor (membrane bound MMP-2 as shown in our study) to be necessary for CAR recognition is not known, an early study identified a low-affinity CLTX receptor which showed high abundance on GBM cell membranes (*76*). Since CD28 has been reported to lower the affinity threshold for TCR activation (*77*), it is possible that interaction between the CLTX tumor-recognition domain and low-affinity GBM receptors requires CD28 co-stimulation to sufficiently initiate CLTX-CAR T cell activation. We noted that along with higher effector function, CLTX-EQ28ζ and CLTX-CD8h28ζ CAR T cells also expressed higher levels of exhaustion markers compared to the other CLTX-CARs evaluated. This observation could result from incomplete activation of the other non-CD28-bearing CLTX-CAR T cells upon tumor binding, as T cell exhaustion markers are induced upon antigen engagement and may also serve as indicators of T cell activation (*78*).

As notes, the composition of the CLTX receptor and its interaction with CLTX peptide is not well defined. The well-characterized inhibition of GBM cell migration and invasion by CLTX (*26, 79*) has been variously attributed to CLTX association with chloride channel CLCN3 and inhibiting the Cl-flux required for cell shape changes during invasion (*27*), or to CLTX inhibiting MMP2 thereby decreasing extracellular matrix cleavage (*26*). Interpretation of immunoprecipitation-based assays of interaction between CLTX and its receptor(s) has been complicated by difficulties distinguishing between direct interaction and indirect association. Here using genetic manipulations we observed that MMP2 expression is required for CLTX-CAR T cell activation, consistent with MMP2 serving as the CLTX receptor or a critical component within a receptor complex. Importantly, CLTX-CAR T cells did not respond to secreted MMP2, thus preserving their activation potential even in a soluble MMP2-rich microenvironment (*80*). The development of CLTX-CAR T cells provide evidence that CAR T cells are able to selectively target membrane-bound form of secreted enzymes, in addition to the previously-reported CAR designs against soluble proteins (*81, 82*). Further, since CAR T cell activation requires direct antigen engagement, our results are consistent with MMP2 being a direct target for CLTX, with their interaction recruiting other associated proteins such as CLCN3 (Fig. 6I and S6H). Similar pattern of CLTX’s association with GBM cells was also suggested by previous studies using CLTX-coated nanoparticles and fusion proteins (*57, 83*). Of note, correlation was observed between MMP2 expression, degranulation (Fig. 6B), *in vivo* CLTX-CAR efficacy (Fig. 6F), and CLTX-Cy5.5 staining intensity (Fig. 6C). Our result suggest that CLTX-Cy5.5 staining of GBM cells could serve as an efficient strategy for identifying the potential responsiveness of GBM tumors to CLTX-CAR T cell therapy.

A generalizable finding arising from our *in vivo* CLTX-CAR studies is the observation that antitumor function of GBM-targeted CARs may be inhibited by adaptive resistance mechanisms of GBM tumors (Fig. 4). Although most of the patient-derived GBM cell lines used in this study did not express PD-L1 (data not shown), induction of PD-L1 expression by GBM xenografts, as well as TS lines, did occur after CAR therapy, and seemed to depend on IFN-γ receptor expression, consistent with the mechanism characterized in metastatic melanomas (*51*). The inhibitory effect of PD-L1 to CLTX-CAR T cells acts mainly on the long-term *in vivo* function, as initial CLTX-CAR activation (degranulation) was not well correlated with the IFNγRA (CD119) expression potentially linked to checkpoint inhibition (Fig. S6D). Our results lead to the potential of correlating tumor IFN-γ receptor expression with CLTX-CAR T cell therapeutic effect, as well as the possibility of combining CLTX-CAR T cells with checkpoint inhibitors for more effective GBM clearance.

Overall, we were able to observe broad GBM-targeting capability of CLTX-CAR T cells, consistent with the wide expression of its receptor, MMP2, in GBM samples (*84*). Of particular importance, CLTX-CAR T cells elicited potent immune responses against tumor cells with no or little expression of other targetable antigens (IL13Rα2, HER2 and EGFR). Through its combination of selectivity and ubiquity, CLTX-CAR T cells address two major hurdles to effective immunotherapy treatment of GBM: reduction of antigen escape while maintaining tumor cell restriction. We suggest that CLTX-CAR T cells are a candidate for clinical development of anti-GBM immunotherapy, circumventing antigen escape either as a single agent, or in combination with other CAR T cells or immunotherapeutic strategies.

## Materials and Methods

### Study design

In this study we evaluated the antitumor potency of CLTX-CAR T cells against GBMs. First, CLTX binding was verified in various GBM samples. All freshly dispersed tumors were processed into single cells following the laboratory’s standard protocol, and PBT-TS lines were generated from freshly dispersed tumor cells and maintained at the standard and well-established condition using neurosphere media. For the experiments on functional evaluation, CAR T cells were generated from three different healthy donors, and tested against multiple GBM models established by TS lines. For *in vitro* assays, 2-5 replicates within each condition were used to sufficiently represent intra-group variations and allow for defining statistical significance. In all *in vivo* experiments, 6-8 week-old NOD/SCID/IL2R-/- (NSG) mice were used, and 4-8 mice were included within each group which enabled us to statistically distinguish tumor sizes and survival rates across groups. Before CAR T cell treatment, mice were grouped with similar average tumor sizes across groups. The health condition of mice was monitored at a daily basis by the Department of Comparative Medicine at City of Hope, with euthanasia applied according to the American Veterinary Medical Association Guidelines. For every single mouse euthanized, brain was collected to confirm the presence of GBM tumors. The pathological conditions of mouse organs were determined by a mouse pathologist from the Veterinary Pathology Program at City of Hope..

### Generation of CLTX-conjugated peptides

Conjugation of CLTX (Alomone Labs) with Cy5.5 fluorescent dye (GE Healthcare) or biotin (Thermo Fisher Scientific) was performed by the Synthetic and Biopolymer Chemistry Core at City of Hope.

### Isolation of primary brain tumor cells, establishment of neurospheres and other cell lines

Primary brain tumor (PBT) cells were obtained from GBM patient resections at COH under protocols approved by the COH Internal Review Board. Resected brain tumor specimens were digested using a human tumor dissociation kit (Miltenyi Biotech Inc) to generate PBT cells. TS lines were subsequently established from PBTs and maintained as described previously (*16, 37*). To generate cells for *in vivo* biophotonic imaging, these cells were engineered to express the ffLuc reporter gene as previously described (*16*). Differentiation of TS lines was performed by withdraw of EGF/FGF in the TS culture media and supplement with 10% fetal calf serum (FCS), as depicted previously (*16*). FB-NSCs were established and characterized as reported in previous studies (*52–54*). Astrocytes and NPCs are differentiated from health donor-derived iPSCs based on established protocols (*85, 86*).

### DNA constructs

All CLTX-CAR constructs contain a CLTX peptide and the cytoplasmic domain of human CD3 zeta, with different spacers including: IgG4EQ [IgG4 with two point mutations (L235E, N297Q) (*7*)]; ΔCH2: IgG4-Fc with the CH2-domain deleted; CD8h: CD8 hinge; L: a synthetic 10 amino acids short linker. CAR constructs also contain CD4 or CD28 transmembrane domains, and CD28 or 4-1BB costimulatory domains. All domains have been previously reported (*43, 45, 87*). A truncated CD19 was also introduced in the construct to allow for potential enrichment and tracking of transduced cells. The firefly luciferase (ffLuc)-GFP construct for tumor biophotonic imaging was generated as described previously (*16*).

### CAR T cell production

Blood products were obtained from healthy donors under protocols approved by the City of Hope (COH) Internal Review Board, and naïve/memory T cell (Tn/mem) isolation followed the procedures described in previous studies (*87, 88*). In brief, peripheral blood mononuclear cells (PBMCs) were isolated by density gradient centrifugation over Ficoll-Paque (GE Healthcare) and then underwent sequential rounds of CliniMACS®/AutoMACS® depletion to remove CD14, CD25-expressing cells, followed by a CD62L positive selection for Tn/mem cells. To generate CAR T cell products, T cells were stimulated with Dynabeads® Human T expander CD3/CD28 (Invitrogen) at a 1:3 ratio (T cell:bead), and transduced with lentivirus to express CAR (MOI=2) in X-VIVO 15 (Lonza) containing 10% FCS with 5 μg/mL protamine sulfate (APP Pharmaceuticals), 50 U/mL rhIL-2 and 0.5 ng/mL rhIL-15. Cultures were then maintained at 37°C, 5% CO_2_ under the same condition of media and cytokines (cytokines were supplied every other day). On day 7 post transduction, the CD3/CD28 Dynabeads were removed from cultures using the DynaMag-50 magnet (Invitrogen). CAR-transduced T cells were enriched by positive selections using anti-CD19 magnetic beads (Stem Cell Technologies). Cultures were propagated for 14-16 days before applying to assays or get cryo-preserved. Mock-transduced T cells were generated by stimulating and culturing Tn/mem cells from the same donors as described above, without lentivirus addition.

### Flow cytometry

Cells were harvested and stained as described previously (*7*). T cell phenotype was detected using fluorochrome-conjugated antibodies against CD3, CD4, CD8 and CD45 (BD Biosciences). Transgene expression was determined by staining for the truncated CD19 (BD Biosciences), and all engineered T cells were gated as the CD3^+^, CD45^+^, CD19^+^ population unless specifically mentioned in figure legends. T cell activation was determined using antibodies against CD69, CD107a, CD137, IFN-γ (BD Biosciences) and PD-1 (BioLegend). Tumor cells were stained with Cy5.5-CLTX, or antibodies against IL13Rα2, HER2, EGFR (BioLegend), CD44 (BD Biosciences) and CD133 (Miltenyi Biotech Inc.). All samples were analyzed via a Macsquant Analyzer (Miltenyi Biotec Inc.) and processed via FlowJo v10.

### In vitro T cell functional assays

To acquire GBM cells for testing CLTX-CAR T cytotoxicity and activity, TSs were dissociated with cold Accutase (Innovative Cell Tec) and resuspended in DMEM:F12 (Irvine Scientific) medium supplied with 10% FCS. CLTX-CAR T cells were then washed and resuspended in the same medium, and added to the PBT cells. To test for degranulation, CLTX-CAR T cells were incubated with GBM cells (E:T=1:1) for 5 h in the presence of CD107a antibody and Golgistop^TM^ protein transport inhibitor (BD Biosciences). To test for CAR T cell killing potency, CLTX-CAR T cells were co-cultured with GBM cells at different E:T ratios for 24-72 h as indicated in individual legends. For the repetitive tumor challenge assay, 4,000 CLTX-CAR T cells were initially co-cultured with 16,000 GBM cells, and then rechallenged with additional 32,000 GBM cells every 48 h for 3 rounds (*48*). Killing was quantified by analyzing viable tumor cells (CD45^−^ DAPI^−^) after co-culture by flow cytometry.

### Immunofluorescence staining

To visualize immunological-synapses, GBM cells were harvested from TS lines as described above, and plated for 12h to adhere. CLTX-CAR T cells were then added to GBM cells and incubated for 3h before fixation with 4% paraformaldehyde (in PBS). Fixed cells were first stained with following antibodies: rabbit-anti-human pCD3ς (Abcam, EP776(2)Y), rabbit-anti-human MMP2 (Abcam, EPR1184) and mouse-anti-human CLCN3 (Abcam, N258/5). Permeabilization and cytoskeleton staining was then performed using an F-actin Visualization Biochem Kit (Cytoskeleton Inc.). To stain for GBM-associated antigens, mouse brains bearing GBM xenografts were harvested, embedded into paraffin and sectioned as described previously (*16*). IF staining was performed on paraffin sections using the procedures of a previous study (*89*), with the following antibodies: goat-anti-human IL13Rα2 (R&D systems, polyclonal), mouse-anti-human EGFR (Dako, DAK-H1-WT), and CLTX-biotin. All slides for immunofluorescence assays were observed with an LSM Airyscan 880 (Zeiss) and images were processed with Zen (Zeiss). Quantification of immunofluoresence images was performed using Image Pro (Media Cybernetics).

### GBM xenograft studies

All mouse experiments were approved by the COH Institutional Animal Care and Use Committee. Orthotopic GBM models were generated using NSG mice as previously described (*50*). Briefly, on day 0, ffLuc^+^ GBM cells (1×10^5^) were stereotactically implanted into the right forebrain. After 8 days, mice were then treated intracranially with 0.5×10^6^ CAR T cells. Tumor volumes were determined by *in vivo* non-invasive optical biophotonic imaging using a Xenogen IVIS 100 as previously described (*16*). For subcutaneous tumor xenografts, GBM cells (5×10^6^) were injected into the left flank of NSG mice and tumors were allowed to grow for 7-14 days. CAR T cells (2×10^6^) were injected intratumorally and tumors sizes were monitored until animal euthanasia when tumor sizes reached 15mm×15mm. To acquire single cells for flow cytometric analysis, xenograft tumors were cut into pieces, physically dissociated and filtered. The pathological conditions of mouse organs were determined by the Veterinary Pathology Program at City of Hope.

### Immunohistochemistry

Mouse brain harvesting, sectioning and immunohistochemistry (IHC) assays were performed as described previously (*16*). Antibodies used in IHC assays include: mouse-anti-human CD3 (Dako, F7.2.38), rabbit-anti-human Granzyme B (eBiosciences, 496B), rabbit-anti-human-PDL1 (Cell Signaling Technology, E1L3N). The slides for IHC assays were scanned via a NanoZoomer 2.0-HT Digital slide scanner (Hamamatsu) and processed with NDP.view2 (Hamamatsu).

### Quantitative real-time PCR

Total mRNA from tumor cells was isolated by RNeasy Mini Kit (Qiagen Inc.). cDNA was then synthesized using an Omniscript RT Kit (Qiagen Inc.). Quantitative real-time PCR was performed using SYBR Green PCR Master mix (Applied Biosystems) in a ViiA^TM^-7 RT-PCR system (Thermo Fisher Scientific). Primer sequences are available upon request. The comparative Ct values of genes of interest were normalized to the Ct value of β-actin. Then, the 2^−Δct^ method was used to determine the relative expression of the genes, while the 2^−ΔΔct^ method was used to calculate the fold changes of gene expression over control.

### Statistics

Data analysis was performed using Prism v6.0 (GraphPad Software) and presented as stated in individual figure legends. Comparisons were determined using Student’s t-test (two groups) or one-way ANOVA (three or more groups). For comparisons between three or more groups, Bonferroni’s Multiple Comparison Tests were used to compare all or selected pairs of data (95% confidence intervals). Comparison of Kaplan-Meier survival data was performed using the Log-rank (Mantel-Cox) test. Detailed comparisons in each experiment are described in figure legends.

### Study Approval

All mouse experiments were approved by the City of Hope Institute Animal Care and Use Committee, Duarte, CA. Use of all human subjects materials (human CAR T cell production and patient-derived GBM spheres) was approved by the City of Hope Institutional Review Board, Duarte, CA.

## Supplementary Materials

### Materials and Methods

Fig. S1. Antigen expression on PBT-TS lines.

Fig. S2. Activation of CLTX-CAR T cells after GBM stimulation.

Fig. S3. The CLTX-EQ-28ζ and CLTX-CD8h-28ζ CARs mediates potent effector function as compared with other CLTX-CAR constructs.

Fig. S4. CLTX-CAR T cells target GBM xenografts.

Fig. S5. CLTX-CAR T cells do not exhibit off-tumor targeting in tumor-bearing mice.

Fig. S6. MMP2 is necessary for CLTX-CAR T cell activation.

Video.S1. Killing of PBT003-4-TS-derived GBMs during a 72h co-culture with the mixture of IL13Rα2-, HER2- and EGFRvIII-targeted CAR T cells

Video.S2. Killing of PBT003-4-TS-derived GBMs during a 72h co-culture with CLTX-EQ-28ζ CAR T cells

## Acknowledgements

We thank the Department of Comparative Medicine, and the cores of Synthetic and Biopolymer Chemistry, Small Animal Imaging, Light Microscopy, Mouse Pathology, Solid Tumor Pathology; as well as Brenda Chang, Juan Ruiz-Delgado and Dr. Lihong Weng for their technical assistance. We thank Dr. James M. Olson for interactive discussions and intellectual feedback on this work.

## Author contributions

Designing research studies: D.W., S.J.F., M.E.B. and C.E.B.; Conducting experiments: D.W., R.S., W.C.C., B.A., S.W., X.Y., A.B., and A.S.; Data acquisition: D.W., R.S. and B.A.; Analysis and interpretation: D.W., D.A., M.E.B. and C.E.B.; Resources: M.G., K.A., Y.S., B.B., C.E.B. and S.J.F.; Writing manuscript: D.W., D.A., J.R.O., M.E.B. and C.E.B.; Supervision: S.J.F., M.E.B. and C.E.B.

## Data and materials availability

PBT-TS lines are available from C.E.B. under a material transfer agreement with City of Hope.

