## Supplementary material for "Chlorotoxin Redirects Chimeric Antigen Receptor T Cells for Specific and Effective Targeting of Glioblastoma": Fig. S1-S6

**Supplementary Materials**

**Materials and Methods**

*Real-time imaging*

For real-time tracing of *in vitro* CAR T cell killing, GBM and CAR T cells were co-cultured and visualized for 72h continuously via an Observer Z1 Live Cell (Zeiss) that contains an incubator with constant 37° C and 5% $CO_2$. Images were processed and movies generated with Zen (Zeiss).

*Detection of cytokine and MMP2 secretion*

To measure the quantity of secreted cytokines, T cells were co-cultured with GBM cells (10,000 $CAR^+$ T cells, 20,000 tumor cells) for 24h. Supernatants were then collected and cytokines were quantified using LEGEND MAX™ Human IFN-γ ELISA Kits (BioLegend). For the detection of MMP2 secretion, GBM cells were cultured (20,000 tumor cells in 200µL of media) for 24 hours and media was harvested. MMP2 secretion was determined using a Human Magnetic Luminex Performance Assay Base Kit, MMP Panel (R&D Systems).

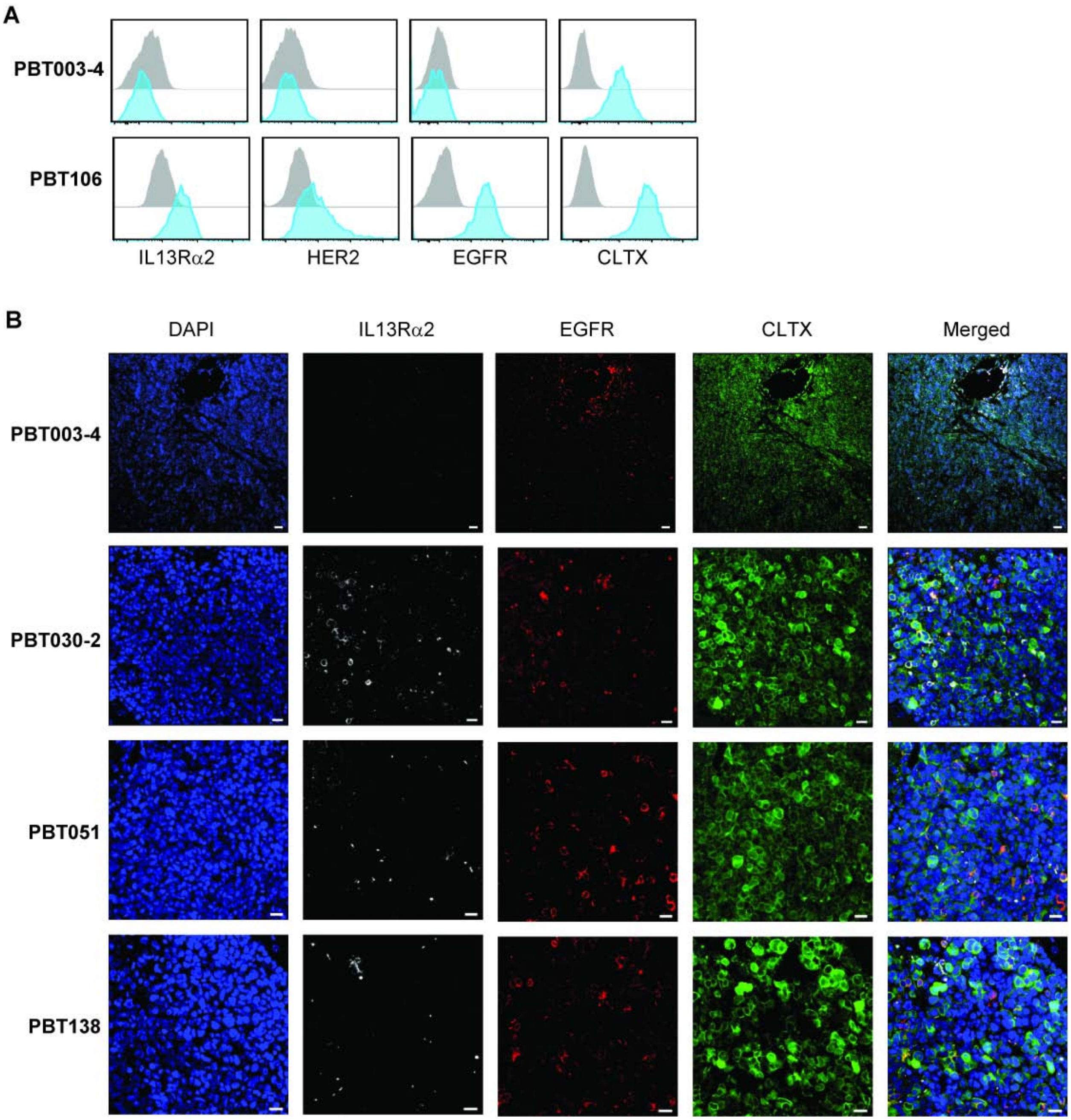


**Fig. S1. Antigen expression on PBT-TS lines.** (**A**) Representative immunostaining of IL13Rα2, HER2 or EGFR, and binding by CLTX-Cy5.5 on GBM cells dissociated from two PBT-TS lines. (**B**) Representative immunofluorescent phenotypes of GBM xenografts established by stereotactically injecting 1 x $10^5$ GBM cells from different PBT-TS lines into the right forebrain of NSG mice. Tumor-bearing mouse brains were harvested 80-100 days after cell injection, and frozen sections were stained with anti-IL13Rα2 or anti-EGFR, or CLTX-biotin, followed by secondary antibodies and then DAPI to identify nuclei. Scale bars: 20 µM. All PBT numbers indicate TS lines.

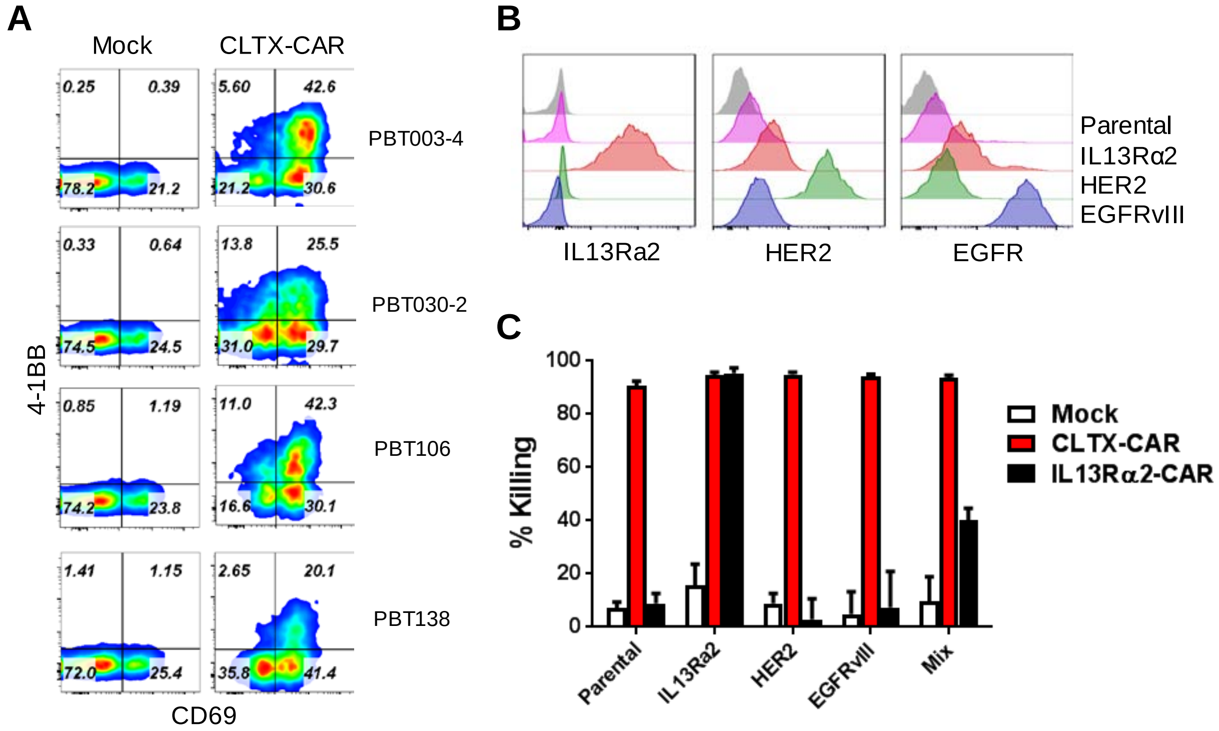


**Fig. S2. Activation of CLTX-CAR T cells after GBM stimulation.** (**A**) Expression of T cell activation markers CD69 and 4-1BB on mock and CLTX-CAR T cells measured by flow cytometry following 24 hours of co-culture with GBM cells from four different TS lines. (**B**) Antigen expression of PBT003-4-TS cells after lentiviral transduction to overexpress IL13Rα2, HER2 or EGFRvIII. (**C**) Cytotoxicity over 48 h co-culture of CLTX- or IL13Rα2-CAR T cells tested against PBT003-4 GBM cells over-expressing IL13Rα2, HER2 or EGFRvIII, or a mixture of PBT003-4 cells (Parental:IL13Rα2:HER2:EGFRvIII = 1:1:1:1). All PBT numbers indicate TS lines.

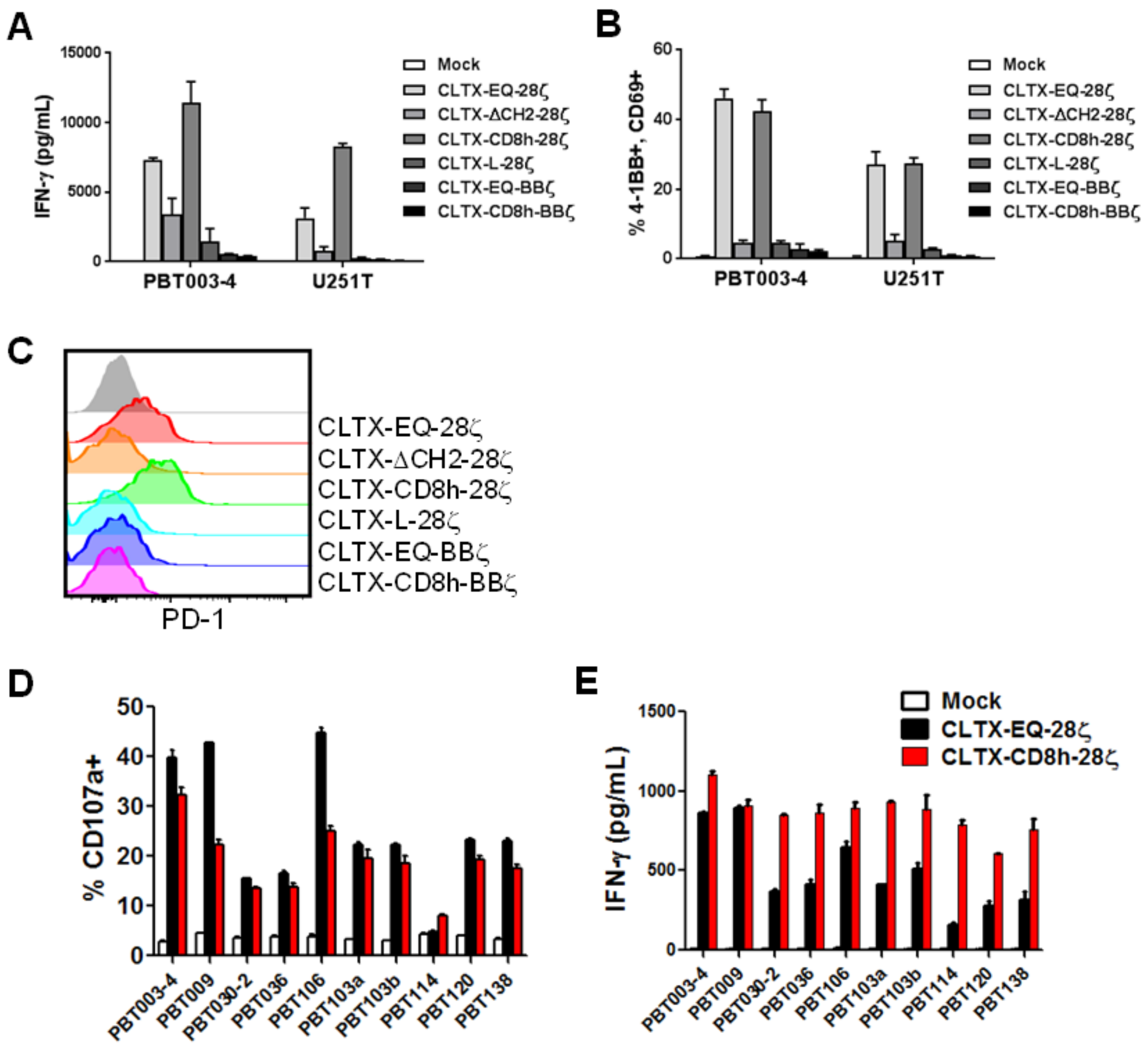


**Fig. S3. The CLTX-EQ-28ζ and CLTX-CD8h-28ζ CARs mediates potent effector function as compared with other CLTX-CAR constructs.** (**A-B**) T cells transduced with different CLTX-CAR constructs were co-cultured with PBT003-4-TS or U251T GBM cells for 24 h. Conditioned media were harvested and secreted IFNγ was measured (**A**), while, in parallel, CAR T cells were harvested for detection of activation markers CD69 and 4-1BB (**B**). (**C**) Histograms of PD-1 surface expression on T cells transduced with different CLTX-CAR constructs and co-cultured with PBT003-4 GBM cells for 24 h. (**D-E**) Degranulation (% CD107a+; **D**) and IFNγ production (**E**) by CLTX-EQ28ζ and CLTX-CD8h28ζ CAR T cells against a panel of GBM cells from 10 PBT-TS lines. All PBT numbers indicate TS lines.

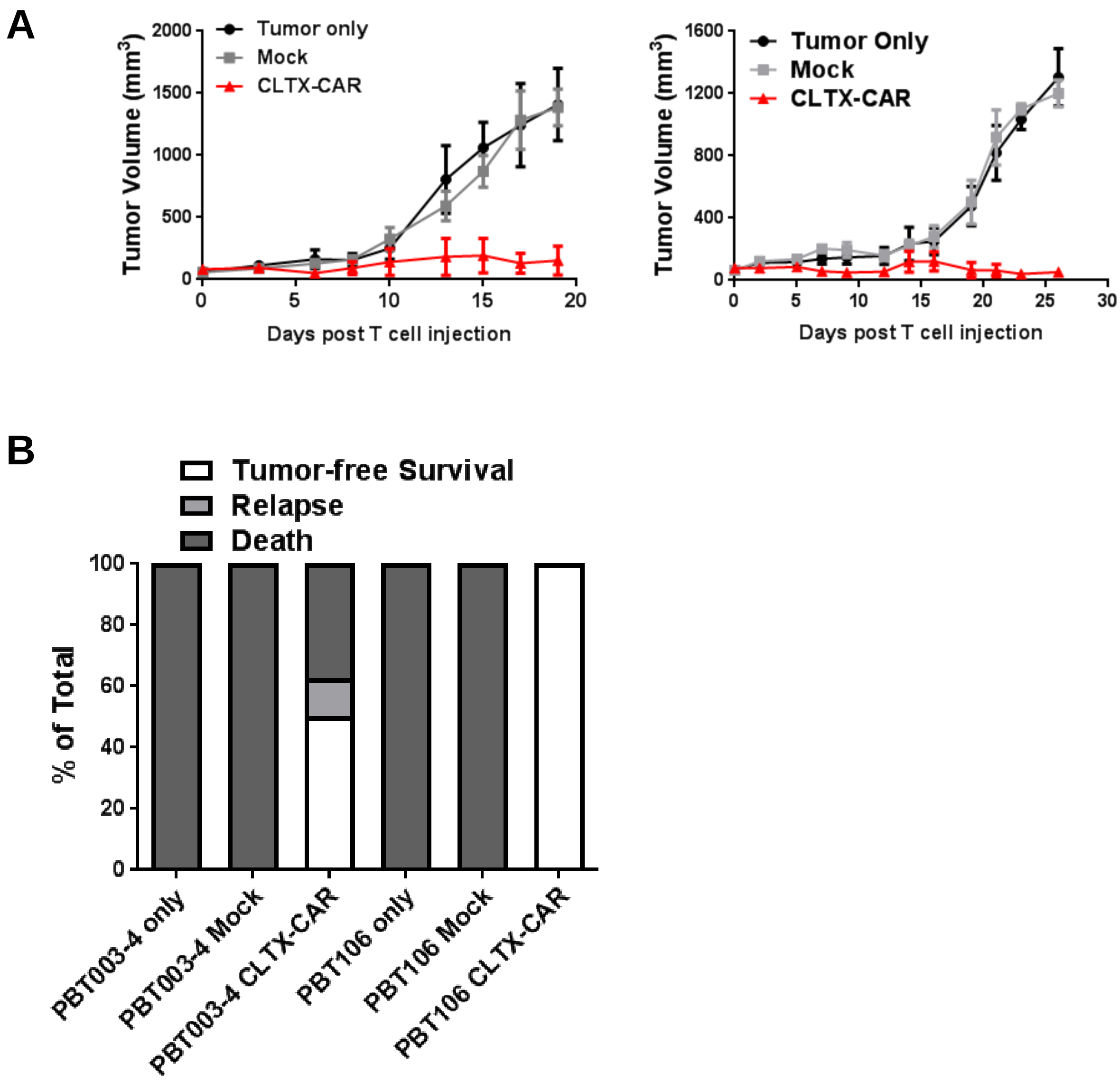


**Fig. S4. CLTX-CAR T cells targeting of GBM xenografts.** (**A**) $5 \times 10^6$ GBM cells from PBT003-4-TS (*left*) or PBT106-TS (*right*) lines were implanted into the right flank of NSG mice. At day 14 post tumor implantation (tumor diameter ~5 mm), $3 \times 10^6$ CLTX-EQ28ζ CAR T cells or mock T cells were injected into the tumors, and tumor size was monitored until tumor diameter reached ~15 mm. (**B**) NSG mice bearing orthotopic PBT003-4-TS or PBT106-TS GBMs received mock or CLTX-CAR T cell treatment as shown in Figure 4. At the end point of the study (D175 post T cell injection), mice were categorized by biophotonic imaging as tumor-free, relapsed or dead (euthanized due to progressive disease). All PBT numbers indicate TS lines**.**

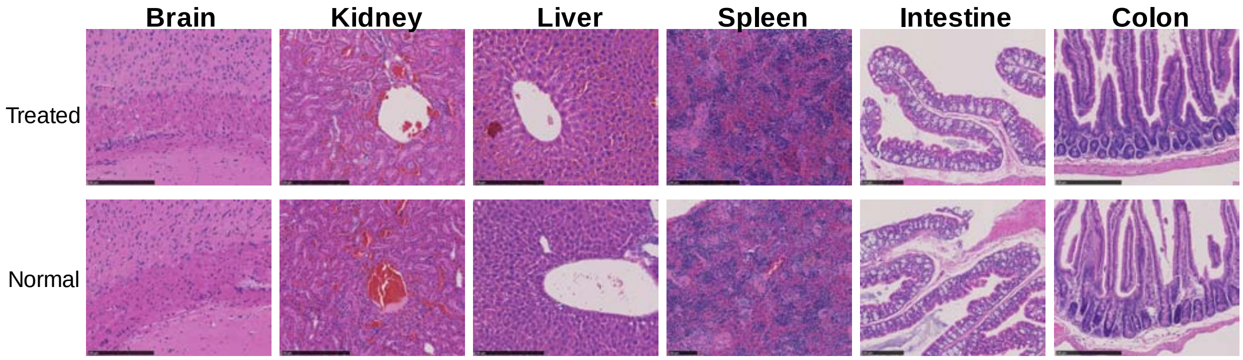


**Fig. S5. CLTX-CAR T cells do not elicit off-tumor pathologies in tumor-bearing mice.** (**A**) NSG mice bearing orthotopic PBT003-4-TS GBM tumors were treated with intracranialy administered CLTX-EQ28ζ CAR T cells. 14 days after T cell administration, organs were harvested for IHC staining and examined for signs of pathological alterations (*upper row*), as compared to the histology of normal organs (*lower row*).

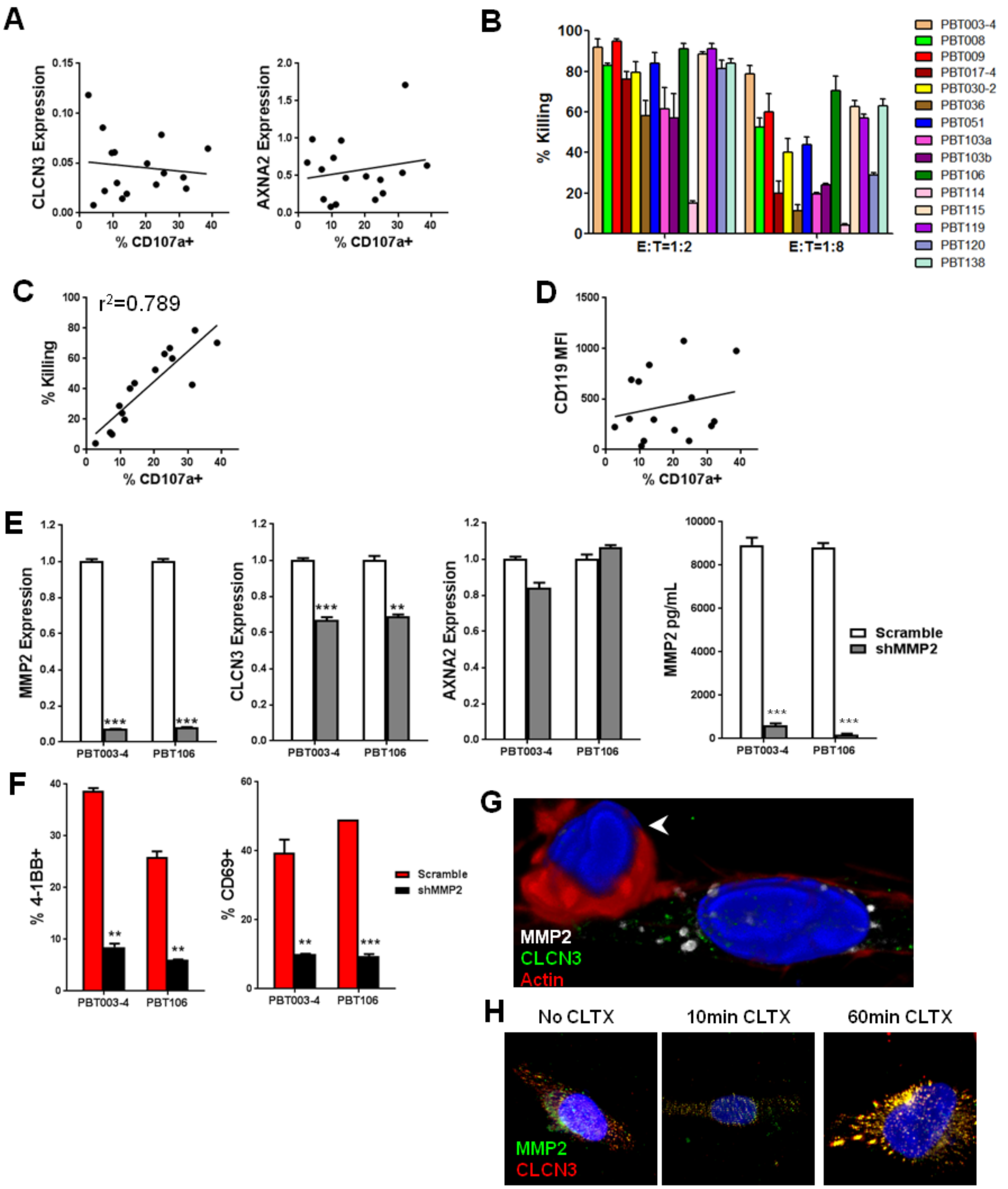


**Fig. S6. MMP2 is necessary for CLTX-CAR T cell activation.** (**A**) Correlation of CLTX-EQ28ζ CAR T cell degranulation (%CD107a+) of CLTX-EQ28ζ CAR T cells with cytotoxicity (% killing) against different GBM-TS lines at E:T=1:8 (96 h). (**B**) Cytotoxicity of CLTX-EQ28ζ CAR T cells tested against

GBM cells from a panel of 15 PBT-TS lines at E:T ratios of 1:2 (over 48 hours) or 1:8 (over 96 hours). (**C**) Relations between CLCN3 (left) and ANXA2 (right) mRNA expression with CLTX-CAR T cell degranulation. (**D**) Relation between CD119 (IFNγRA) staining intensity (MFI) on target cells and CLTX-CAR T cell degranulation. (**E**) MMP2, CLCN3 and AXNA2 mRNA expression (left three panels), and soluble MMP2 levels (right panel), for PBT003-4 and PBT106 GBM cells after lentiviral transduction of shMMP2 RNA or scramble shRNA. (**F**) Expression of activation markers CD69 and 4-1BB by CLTX-EQ28ζ CAR T cells after 24 h co-culture with PBT003-4-TS or PBT106-TS GBM cells transduced with shMMP2 RNA or scrambled control shRNA. (**G**) PBT106-TS GBM cells co-cultured with IL13Rα2-CAR T cells for 2 h, and stained for MMP2, CLCN3 and actin polarization (immunological synapse). Arrowhead: T cell. (**H**) PBT003-4-TS GBM cells were exposed to CLTX peptide for 10 or 60 min, and then stained for co-localization of MMP2 and CLCN3. **$p<0.01$, ***$p<0.001$ using a unpaired student's t test. All PBT numbers indicate TS lines.
